# Hydroxyzine inhibits SARS-CoV-2 Spike protein binding to ACE2 in a qualitative *in vitro* assay

**DOI:** 10.1101/2021.01.04.424792

**Authors:** Maria Dolores Rivas, Jose Maria Rafael Saponi-Cortes, Jose Zamorano

## Abstract

COVID-19 currently represents a major public health problem. Multiple efforts are being performed to control this disease. Vaccinations are already in progress. However, no effective treatments have been found so far. The disease is caused by the SARS-CoV-2 coronavirus that through the Spike protein interacts with its cell surface receptor ACE2 to enter into the host cells. Therefore, compounds able to block this interaction may help to stop disease progression. In this study, we have analyzed the effect of compounds reported to interact and modify the activity of ACE2 on the binding of the Spike protein. Among the compounds tested, we found that hydroxyzine could inhibit the binding of the receptor-binding domain of Spike protein to ACE2 in a qualitative *in vitro* assay. This finding supports the reported clinical data showing the benefits of hydroxyzine on COVID-19 patients, raising the need for further investigation into its effectiveness in the treatment of COVID-19 given its well-characterized medical properties and affordable cost.

## Introduction

Coronavirus Disease 2019 (COVID-19) has emerged as the most severe pandemic disease over the last decades (1). It has become a major public problem, collapsing health care systems worldwide. Multiple efforts are being addressed to develop strategies to control this disease. Thus, vaccines have recently been approved and vaccinations have started (2). In contrast, although several potential treatments are being investigated and even approved, their efficacy remains limited (3,4). Therefore, continuing the investigation of COVID-19 treatments remains of great importance for the management of this disease.

COVID-19 is caused by the SARS-CoV-2 coronavirus (1). Cellular infection is facilitated by the binding of the receptor-binding domain (RBD) of the viral Spike protein to the angiotensin-converting enzyme 2 (ACE2), its receptor on the cell surface. Therefore, compounds able to block the interaction of Spike with ACE2 may be useful in preventing virus progression at the early stages of infection. In this regard, Kulemina and Ostrov reported in 2011 the existence of several small known compounds that could interact with ACE2 modifying its *in vitro* activity (5). Among them, hydroxyzine was found to have a statistically significant effect on ACE2 efficiency. They proposed that the binding of compounds to ACE2 could provoke conformational changes that would affect angiotensin II binding. We hypothesized that the binding of hydroxyzine to ACE2 might also interfere with SARS-CoV-2 binding to ACE2. This idea may help to understand the reported beneficial effect of hydroxyzine in this disease (6,7). To investigate it, we analyzed the effect of hydroxyzine on an *in vitro* assay that qualitatively determines the binding of recombinant Spike to ACE2.

## Materials and Methods

Drugs: hydroxyzine, labetalol, and other compounds were purchased from Sigma-Aldrich (St. Louis, MO).

### Binding assays

The RayBio COVID-19 spike-ACE2 binding assay (Cat: Cov-ACE2S2) was obtained from RayBiotech, Inc. (Peachtree Corners, GA). This assay is a qualitative method to characterize the binding of SARS-CoV-2 Spike receptor-binding domain (RBD) to ACE2 in the presence of potential inhibitors The Binding assay experiments were performed as indicated in the instruction manual. The binding of RBD to ACE2 was determined in a Tescan spectrophotometer at 450 nm.

## Results

We analyzed the effect of selected compounds on the binding of recombinant Spike RBD protein to ACE2. To this end, the indicated amounts of drugs were mixed with RBD and added to wells containing immobilized ACE2 protein. We found that hydroxyzine inhibited in a dose-dependent manner the binding of RBD to ACE2 protein (Figure 1). In contrast, labetalol, another drug reported to interact with ACE2 (5), did not affect RBD/ACE2 interaction.

**Figure 1.**
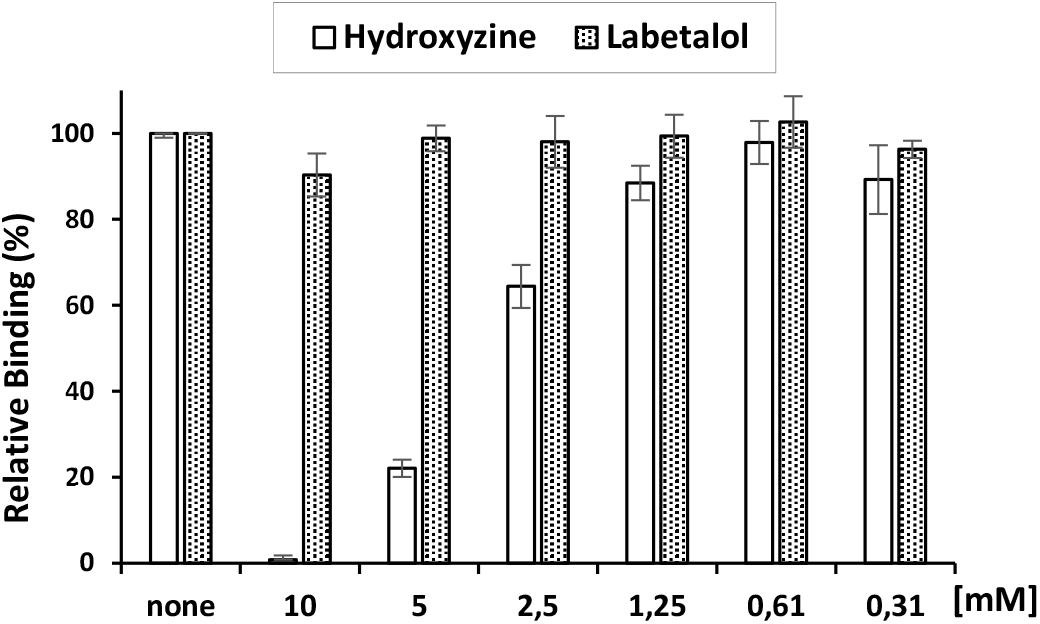
Hydroxyzine competes with Spike RBD protein to bind ACE2. The indicated amounts of Hydroxyzine (empty bars) and Labetalol (dotted bars) were incubated with recombinant spike RBD protein and added to a plate containing recombinant ACE2 as indicated in RayBiotec protocol. This is representative of three different experiments.

These results suggest that hydroxyzine could compete and interfere with the binding of Spike RBD to ACE2. This was further supported by the fact that preincubation of ACE2 with hydroxyzine inhibited to a greater extent the posterior binding of RBD to ACE2 (Figure 2).

**Figure 2.**
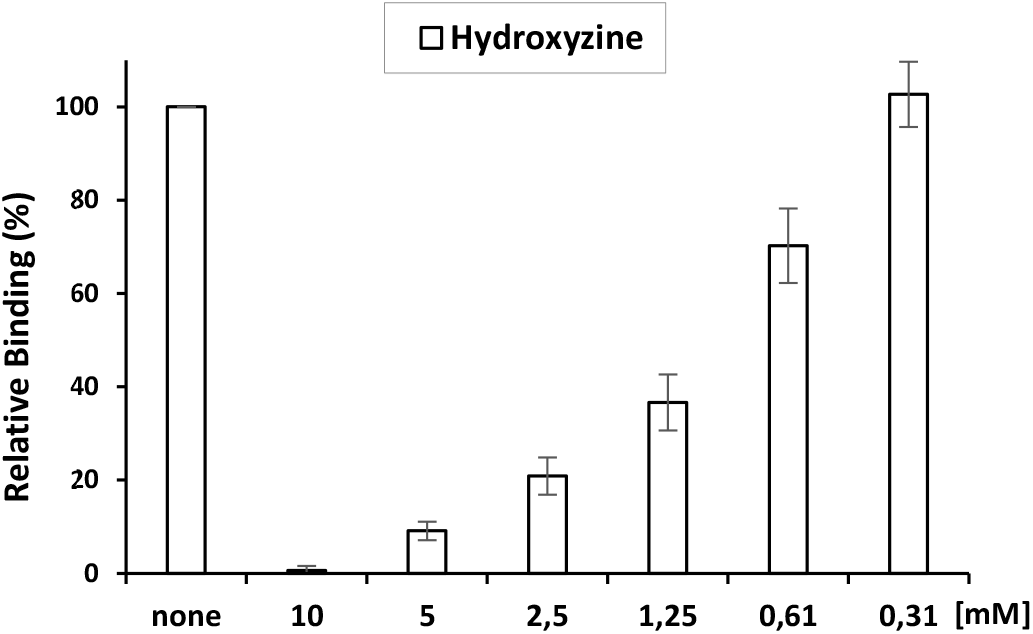
Hydroxyzine inhibits the binding of Spike RBD protein to bind ACE2. A plate containing recombinant ACE2 was preincubated with the indicated amount of hydroxyzine overnight. Then, hydroxyzine was washed out and the binding of spike RBD to ACE2 was determined as indicated in RayBiotec protocol. This is representative of three different experiments.

## Discussion

In this study, we have found that hydroxyzine competed and interfered with viral Spike protein to bind ACE2 in an *in vitro* qualitative assay. These findings are in agreement with the reported *in vitro* antiviral activity of hydroxyzine (6). Hydroxyzine and other antihistaminic drugs have already been shown to have some beneficial effects in treating COVID-19 (6,7). Thus, hydroxyzine has been reported to reduce the incidence of SARS-CoV-2 infection in older people and the mortality in hospitalized patients (6,7). Although these effects may be explained in part by its antihistaminic properties (7), our findings suggest an additional potential effect by blocking the binding of Spike to ACE2. Hydroxyzine is one of the best known and prescribed drugs among the first-generation antihistamines. Short-term use is generally well tolerated, with the most frequent side effects being drowsiness and dry mouth. Taken together the mentioned evidences, the efficacy of hydroxyzine as a treatment for COVID-19 would be worth further investigation. Considering also that it is an affordable and well-characterized drug, it would allow access to a large number of patients worldwide who may not have access to new and more expensive drugs in development.

